# Genomics of PDGFR-rearranged hypereosinophilic syndrome

**DOI:** 10.1101/2022.09.19.508443

**Authors:** Esther Rheinbay, Meifang Qi, Juliette M. Bouyssou, Andrew J. Oler, Lauren Thumm, Michelle Makiya, Irina Maric, Amy D. Klion, Andrew A. Lane

## Abstract

A subset of patients with hypereosinophilia have a clonal hematologic neoplasm. These neoplastic hypereosinophilic syndromes (HES) can harbor somatic DNA rearrangements involving a PDGF receptor gene, *PDGFRA* or *PGDFRB*, most frequently *FIP1L1::PDGFRA*. Patients with PDGFR*-*rearranged HES are overwhelmingly males, and the hypereosinophilia responds robustly to the multi-kinase inhibitor imatinib. Unbiased genomics of PDGFR-rearranged HES have not been comprehensively studied. Here, we compared whole genome sequencing of eosinophil DNA from patients with PDGFR-rearranged HES at diagnosis (tumor) versus matched “normal” DNA. Other than the PDGFR rearrangement itself, we detected no recurrent coding somatic point mutations, insertions/deletions, or DNA copy number alterations in the neoplastic eosinophils; instead, there were additional somatic events private to each tumor. The tumors had a linear clonal structure with the PDGFR rearrangement as the initial driver event. We mapped the breakpoints of a novel *IQGAP2::PDGFRB::UVRAG* intra- and inter-chromosomal fusion event in one patient. Non-coding analysis found no recurrent abnormalities, including in eosinophilic leukemia-associated promoters or enhancers. The sex chromosomes showed no recurrent alterations to provide an obvious explanation for the disease’s extreme male bias. We conclude that neoplastic HES has relatively simple genomics and that the only unambiguous recurrent driver event is the PDGFR rearrangement itself.

**Key Points:**

- PDGFR-rearranged hypereosinophilic neoplasms have simple, linear genetic structure with the rearrangement as the likely initiating event
- There are no obvious recurrent additional or sex-biased genomic events that cooperate with PDGFR rearrangement to drive the neoplasm

## Introduction

The World Health Organization classification of myeloid neoplasms defines a category as Myeloid/Lymphoid Neoplasms Associated with Eosinophilia and Rearrangements of *PDGFRA, PDGFRB*, or *FGFR1*, or *PCM::JAK2*^1^. The most frequent neoplasms in this category are associated with *FIP1L1::PDGFRA* (F/P) fusions, created by a somatic, acquired interstitial deletion on chromosome 4q12^2–5^. Patients with F/P or rearrangements of *PDGFRB* typically present with hypereosinophilic syndrome (HES), defined as >1.5×10^6^ eosinophils/mL blood and evidence of eosinophil-mediated end organ manifestations^6^.

PDGFR-rearranged hypereosinophilic syndrome (HES) has an overwhelming male predominance (>90%) that is not understood and not observed in other HES^2–4^. PDGFR-associated HES is also exquisitely sensitive to the tyrosine kinase inhibitor imatinib, which targets the constitutive PDGF receptor-signaling driven by the fusions^1,5^. Our goal was to investigate PDGFR-associated HES genomics to better understand these features.

## Methods

Purified eosinophils and peripheral blood mononuclear cells (PBMCs) were isolated from whole blood of patients with HES and a PDGFR abnormality. DNA was prepared for whole-genome sequencing (WGS) using a “PCR-free” library protocol. Sequencing analyses were performed in the TERRA (Terra.bio) platform using established workflows^7^, with custom local analysis in Python. Details in **Supplementary Methods.**

## Results and Discussion

We performed whole-genome sequencing (WGS) on eosinophils (‘tumor’) from 11 patients with PDGFR-related HES (**Table S1**). Paired ‘normal’ samples were PBMCs collected in hematologic and molecular remission after imatinib treatment, in the setting of normal eosinophil counts and no detectable PDGFR rearrangement (**Figure 1A; Supplementary Figure 1A**). The median age at tumor sampling was 43 (range 13-60; **Table S1**); there were 10 males and one female (expected sex bias^3^). Nine males had evidence of a *FIP1L1::PDGFRA* fusion. One female had a *PDGFRB::ETV6* rearrangement, and one male had a complex rearrangement involving *PDGFRB* (see below). Mean sequencing depth was 83 (range 61-144) for tumor and 91 (range 67-125) for normal (**Supplementary Figure 1B; Table S2**).

**Figure 1.**
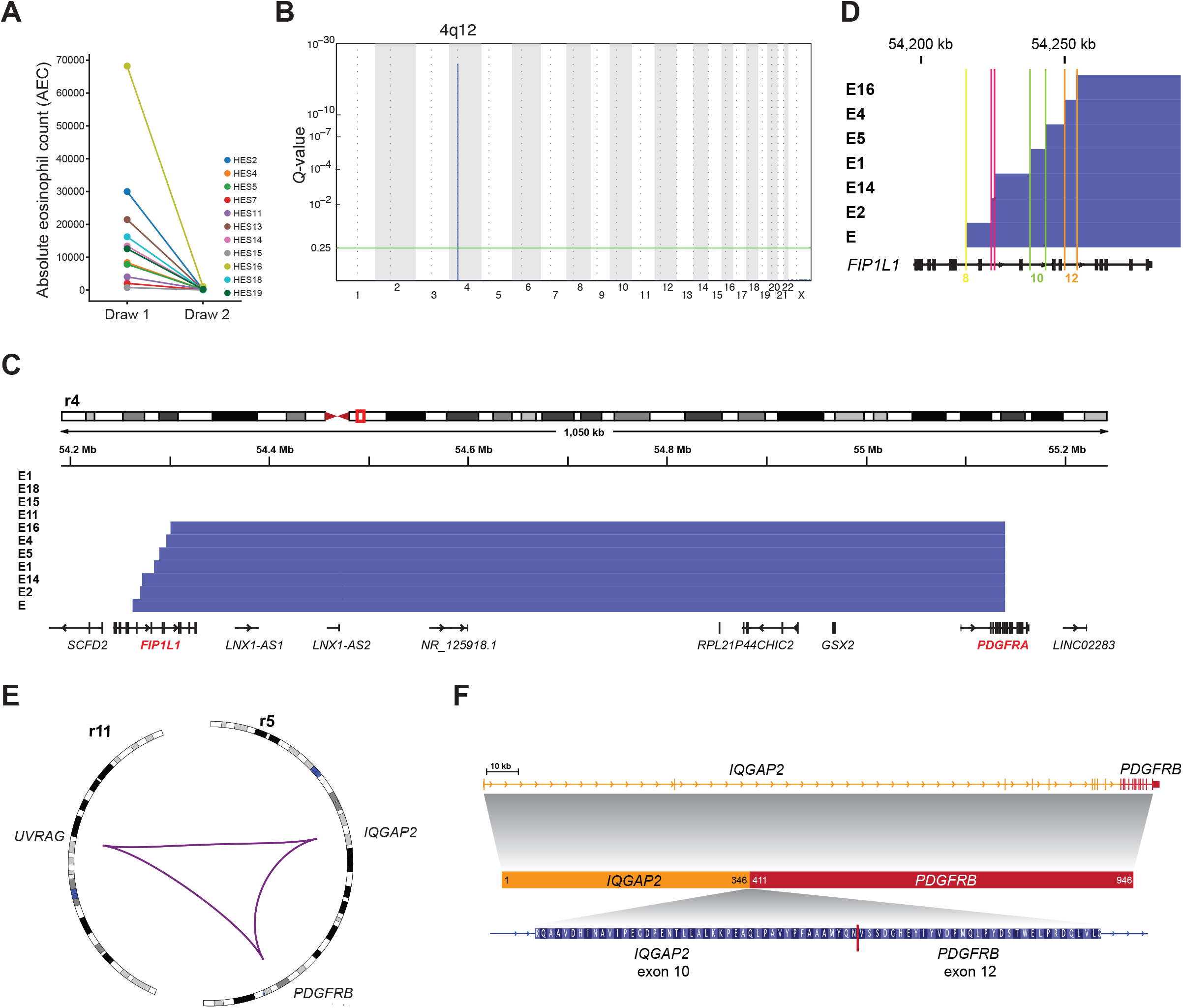
Genomic characterization of PDGFR rearrangements. **A,** Absolute eosinophil count (AEC) before (tumor) and after (normal) treatment with imatinib. **B**, GISTIC2 plot depicting the significantly recurrent interstitial deletion on chromosome 4. **C**, Genomic view of the *FIP1L1::PDGFRA* locus showing detected deletions (blue bars) in most samples. **D**, Detailed view of the breakpoints inside the *FIP1L1* gene. Colored lines show location inside introns labeled with matching intron numbers (based on transcript NM_001134937.1). **E,** Circos^20^ plot of the complex *PDGFRB* rearrangement in patient HES11. Purple lines indicate genomic fusions between chromosomal regions. **F**, Gene locus and inferred protein fusion between *PDGFRB* and *IQGAP2*.

The copy number landscape of F/P HES was “quiet,” with the interstitial deletion between *FIP1L1* and *PDGFRA* as the only recurrent event (**Figure 1B**). *FIP1L1::PDGFRA* was also the most frequently identified structural variation (SV) (**Table S1**). No evidence of F/P was detected in paired normals. All breakpoints in *PDGFRA* occurred within exon 12 (**Figure 1C**). *FIP1L1* breakpoints spanned a ~45-kb region covering introns 8-12 (**Figure 1D; Table S1**).

SV analysis confirmed *PDGFRB* fusions in two tumors with *PDGFRB* events by FISH (**Table S1**). One was an *ETV6::PDGFRB* fusion, reported in HES^8^. The other comprises a previously unreported rearrangement involving *IQGAP2* and *PDGFRB* on chromosome 5 and *UVRAG* on chromosome 11. This SV created an in-frame protein fusion between exon 10 of *IQGAP2* and exon 12 of *PDGFRB*, with the other ends of these breakpoints connecting to the first intron of *UVRAG* (**Figure 1E-F; Supplementary Figure 1C**).

Single-nucleotide variants (SNVs), insertions and deletions (indels) occurred with a median rate of 0.37 events per megabase (range 0.23 to 0.58), comparable to myeloproliferative neoplasms (Myeloid-MPN: 0.38 mutations/megabase^9^) and lower than other hematologic malignancies (**Figure 2A**). The absolute number of SNVs and indels in HES tumors ranged from 653 to 1652 (**Figure 2B; Supplementary Figure 2A**).

**Figure 2.**
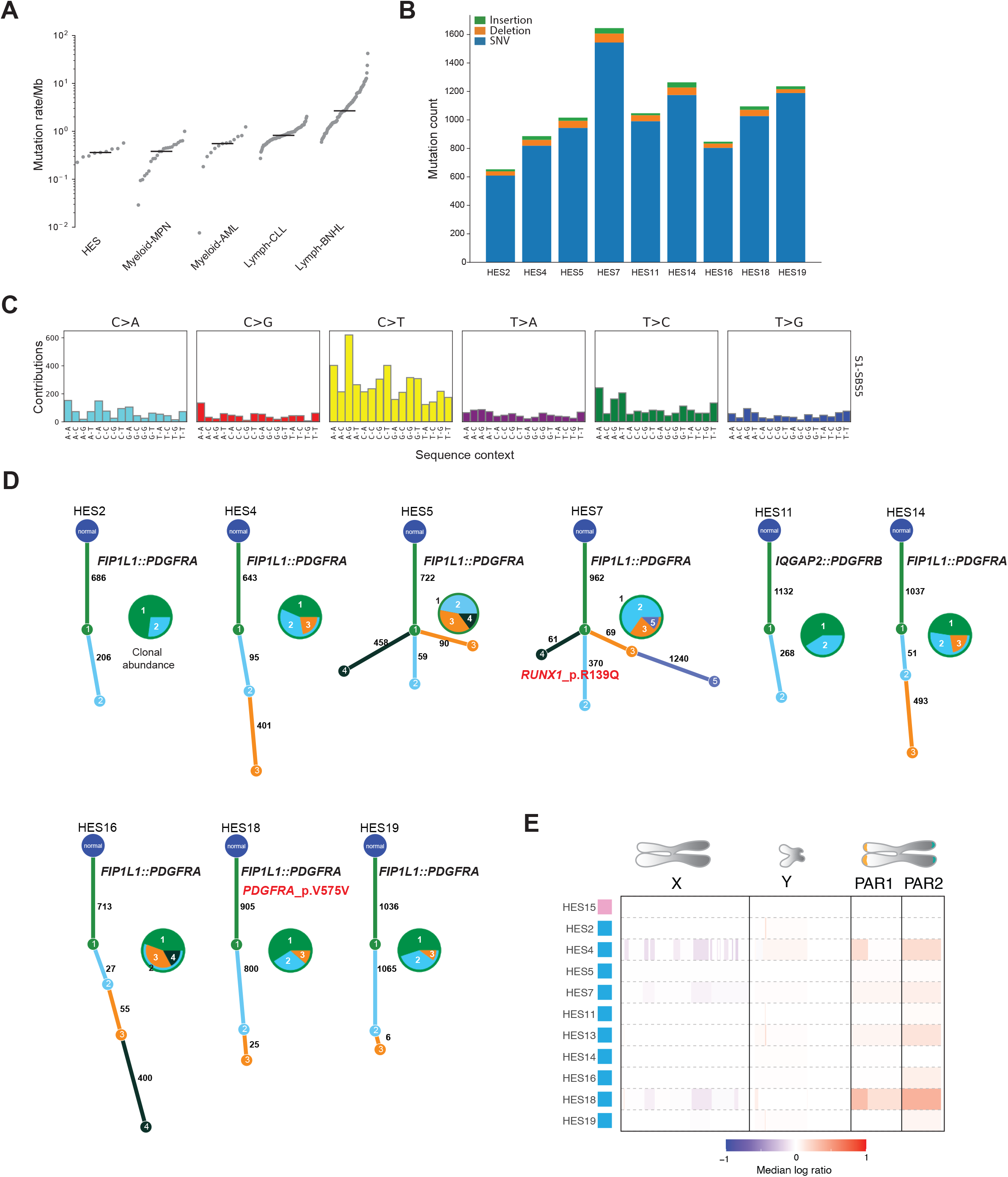
Genomic analyses of PDGFR-rearranged HES. **A,** Mutation rate comparison between HES and other myeloid malignancies. MPN: Myeloid proliferative neoplasm; AML: Acute myeloid leukemia. CLL: Chronic lymphocytic leukemia. BNHL: B-cell non-Hodgkin lymphoma^9^. Each gray dot indicates a sample. **B**, Number of single-nucleotide variants (SNVs; blue), deletions (orange) and insertions (green). **C**, Mutational signature profile for HES samples. The height of each bar indicates the number of mutations in each context (on the x-axis) and base change type (distinguished by color). **D,** Inferred phylogenies for single tumors. Pie charts show the abundance of each clone in each sample. Numbers inside pies correspond to clone ID numbers on the tree. Numbers on tree branches indicate the number of mutations in each branch. Protein-coding alterations in known cancer genes are marked in red. **E**, Copy number calls as median log ratio (tumor/normal) for the sex chromosomes X and Y and two pseudo-autosomal regions (PARs). Pink box: female; blue box: male.

Signature analysis revealed COSMIC SBS5 (C>T transitions) as the most prominent mutational pattern (cosine similarity 0.77) (**Figure 2C; Supplementary Figure 2B**). SBS5 is found across all tumor types, including myeloid neoplasms^10,11^, and exhibits clock-like behavior that correlates with age. No signature suggesting exposure to environmental (e.g., smoking, ultraviolet light) or endogenous (e.g., APOBEC or AID) mutagens was detected.

We next investigated clonal structure using single-sample inference^12^. Most tumors exhibited linear phylogenies with one or two subclones (**Figure 2D**), with two (HES5 and HES7) having branched evolution. The PDGFR fusion occurred in the truncal clone in all cases. We detected a subclonal missense mutation in *RUNX1* in HES7 causing a Arg139Gln amino acid change, previously observed in MDS^13^ and familial platelet disorder^14^. In total, 87 non-silent somatic mutations in protein-coding genes were called in HES tumors, with no recurrently mutated genes. With the caveat of small cohort size, this suggests that there are no obvious additional somatic driver mutations in HES beyond the PDGFR rearrangement.

Identifying recurrent somatic, non-coding driver mutations is challenging because they may not involve the identical base and there is no exonic structure to guide variant classification^15^. Our WGS depth provided sufficient power to interrogate GC-rich promoters and 5’UTRs underrepresented in older studies (**Supplementary Figure S3**)^15,16^. To focus on potential eosinophil-relevant non-coding regions, we analyzed H3K27 acetylation (active promoters and enhancers) and CTCF (chromatin regulatory hub boundaries) sites from chromatin immunoprecipitation-sequencing (ChIP-seq) in EOL1 eosinophilic leukemia cells, which harbor a *FIP1L1-PDGFRA* fusion. No recurrent mutations (>1 mutation per element) were found in these regions, or in transcription start sites, 3’UTRs, and 5’UTRs of protein-coding genes.

Given that the paired normal samples were peripheral blood mononuclear cells collected from patients at a time of HES remission, we searched for background clonal hematopoiesis (CH) in 159 CH-associated genes^17^. Only one non-silent variant was identified, a *CSF1R* substitution (p.R144C, 7% VAF) in patient HES15. Rare CH detection was limited by the expected low frequency of expanded clones. With a median sequencing depth of 91, we achieved 84% power to detect variants with 5% VAF and 52% power to detect variants with 3% VAF.

Next, we performed a custom sex-chromosome analysis to detect somatic variants that could explain the male bias of PDGFR-associated HES (**Supplementary Methods**). No broad or focal copy number changes targeting specific genes were found on chrX or chrY. Importantly, no clonal loss of the Y chromosome was observed, as previously described in leukocytes of aging men^18^. There were no recurrent point mutations in protein-coding genes or regulatory elements, including in tumor suppressors that escape X-inactivation (EXITS genes) or their homologs on chrY^19^.

Here, we show that PDGFR-rearranged HES has simple genomics. We did not observe any frequent cooperating mutation, including in known cancer genes or in genes associated with age-related clonal hematopoiesis. Most of the tumor samples demonstrated a linear phylogeny, and the PDGFR rearrangement was truncal - part of the founding clone - in all cases. This may explain why patients have robust and sustained response to kinase inhibitors.

There was no obvious genetic feature that explains the extreme male bias of *FIP1L1::PDGFRA*- associated HES. It is possible that hormonal or immune environment differences predispose men, or conversely, protect women, from the disease. Alternatively, dosage imbalances in X or Y genes between men and women^23^, even in the absence of a mutation, may predispose eosinophils to acquire PDGFR rearrangements or exert positive selection on mutant cells. Future work in male and female experimental systems modeling eosinophil transformation may address these questions.

There are important limitations to this work. First, in this rare, orphan disease the sample size was limited despite recruiting from a national eosinophil disorders clinic. This constrains our ability to identify events occurring in a subset of patients, and severely limits identification of germline predisposition. Second, the “normal” controls were remission peripheral blood cells, which could miss a mutation acquired in all hematopoietic cells that predisposes to HES. To address this, we manually curated all variant calls in the normals for genes associated with CH and blood cancer. However, this could miss novel predisposing mutations. These limitations might be overcome in a future study using non-hematopoietic normal tissue as reference.

## Acknowledgments

We thank Chip Stewart, Elizabeth Martin, Ignaty Leshchiner, Jason Nomburg, and Kirsten Kubler for analysis support and Alexis Berry and Sana Mahmood for assistance in preparing the DNA samples.

This work was supported by the Mark Foundation for Cancer Research (AAL) and the Bertarelli Rare Cancers Fund at Harvard Medical School (AAL and ER). AAL is a Scholar of the Leukemia & Lymphoma Society. This work was supported in part by the Division of Intramural Research, NIAID, NIH.

## Authorship Contributions

ER, ADK, AAL: study design, data analysis, manuscript preparation and editing. MQ, JMB, AJO, LT, MM, IM: performed experiments, data analysis, manuscript editing.

## Disclosures

AAL is a consultant for Qiagen. AAL has received research support from AbbVie and Stemline Therapeutics.

## Notes

### Competing Interest Statement

AAL is a consultant for Qiagen. AAL received research support from AbbVie and Stemline Therapeutics.

