## Supplementary text and figures for "Genomics of PDGFR-rearranged hypereosinophilic syndrome"

#### Supplementary Methods

##### Patients and Sample Preparation

All patients were enrolled on an NIAID IRB-approved research protocol that allowed DNA sequencing (NCT00001406) and signed written consent. All cases met WHO criteria for myeloid/lymphoid neoplasms with eosinophilia and recurrent rearrangements of *PDGFRA*, *PDGFRB*, and *FGFR1,* or *PMC1*::*JAK2* and had molecular evidence of a PDGFR abnormality detected by reverse transcriptase polymerase chain reaction (*FIP1L1*::*PDGFRA*) and/or fluorescence in situ hybridization (*FIP1L1*::*PDGFRA* and *PDGFRB* fusions). Granulocytes and peripheral blood mononuclear cells (PBMC) were isolated from whole blood by density gradient centrifugation (Ficoll-Paque Plus; GE Healthcare). After red blood cell lysis, eosinophils (tumor) were purified from the granulocyte layer by negative selection on an AutoMacs using the Eosinophil Purification Kit Purity (Miltenyi Biotech) magnetic bead selection as previously described[1]. Eosinophil purity was >97% for all tumor samples based on cytospin analysis of 300 cells. Cells (eosinophils and PBMC) were snap frozen in aliquots of $10^7$ cells/mL and stored in liquid nitrogen. DNA was prepared from thawed cells using Gentra Puregene Cell Kit (Qiagen) according to the manufacturer's instructions.

##### Construction of PCR-free whole genome sequencing libraries

An aliquot of genomic DNA (350 ng in 50 µL) was used as the input into DNA fragmentation (also known as shearing). Shearing was performed acoustically using a Covaris focused-ultrasonicator, targeting 385 bp fragments. Following fragmentation, additional size selection was performed using SPRI cleanup. Library preparation was performed using a commercially available kit provided by KAPA Biosystems (KAPA Hyper Prep without amplification module, product KK8505), and with palindromic forked adapters with unique 8-base index sequences embedded within the adapter (purchased from Roche). Following sample preparation, libraries were quantified using quantitative PCR (kit purchased from KAPA Biosystems), with probes specific to the ends of the adapters. This assay was automated using Agilent's Bravo liquid handling platform. Based on qPCR quantification, libraries were normalized to 2.2 nM and pooled into 24-plexes.

##### DNA sequencing (NovaSeq 6000)

Sample pools were combined with NovaSeq Cluster Amp Reagents DPX1, DPX2 and DPX3 and loaded into single lanes of a NovaSeq 6000 S4 flow cell using the Hamilton Starlet Liquid Handling system. Cluster amplification and sequencing occurred on NovaSeq 6000 Instruments utilizing sequencing-by-synthesis kits to produce 151 bp paired-end reads. Output from Illumina software was processed by the Picard data-processing pipeline to yield CRAM or BAM files containing demultiplexed, aggregated aligned reads. There was no significant cross-contamination from other individuals[2], or oxoguanine mutation artifact[3].
All sample information tracking was performed by automated LIMS messaging. **Table S2** contains the sequencing metrics.

#### Structural variants

Somatic structural variants (genomic rearrangements, insertions, large deletions) were called with Manta[4] with post-filtering previously described[5], requiring at least 10 split reads or read pairs in the tumor, at most 1 such read in the matched normal sample, and a minimum variant allele fraction of 0.05. *PDGFR* structural variants were manually reviewed for accuracy and to confirm exact breakpoints. FACETs[6] (https://github.com/vanallenlab/facets workflow in TERRA with default parameters) was used to call absolute and allele-specific copy number variants (CNV) from WGS with adjustments for sex chromosome calling (Supplementary Methods).

#### SNVs and indels

Single nucleotide variations (SNVs) were called with MuTect version GATK3 v1.1.6[7], MuTect2.0 version GATK3 "3.6-97-g881c5e9"[8] and Strelka2[9]. Insertions and deletions (indels) were identified with two algorithms, MuTect2 and Strelka2 within the TERRA framework, and only indels identified by both methods were used for downstream analysis. Extensive filtering was performed using a cohort-specific panel-of-normals and a local realignment filter[5]. Purity estimates were obtained with using only SNVs and the ABSOLUTE method[10].

#### Mutational signatures

De novo mutational signature discovery was performed on the filtered mutation call set (see above) with SignatureAnalyzer[11] in TERRA (workflow signatureanalyzer snapshot 8). We discovered a signature remotely resembling COSMIC SBS43 (cosine similarity 0.39), a presumed sequencing artifact (**Supplementary Methods**; **Supplementary Figure 2B-F**). Consistent with a non-biologic etiology, mutation events exhibited extreme strand and read orientation bias, specificity for T-C>A-C (A-T>G-G reverse complement) context and base change and low variant allele fraction, which can be characteristic of artifact mutations[3]. A filter based on the mutational signature that recapitulated these mutations was applied to decrease the mutational noise (a mutation was filtered out if its probability for the associated signature was >0.25). This filter is conservative, as it might remove true variants in samples that are largely dominated by the artifact signature (**Supplementary Figure 2B-F**).

#### Copy number

Copy number segment calls from FACETs were postprocessed by excluding regions overlapping the GATK's CNV_and_centromere_blacklist.hg19.list and human centromere exclusion sites with BedTools subtractBed[12]. Recurrent copy number events were identified with GISTIC2 version 2.0.23[13] including the X chromosome (option -rx 0).

To identify copy number segments on the sex chromosomes and pseudoautosomal regions, we adapted the model underlying FACETs[6], enabling us to estimate copy number for X, Y and the two PAR regions PAR1 and PAR2 separately (**Methods**; **Supplementary Figure S3**). This approach required treating the PARs in X and Y as two separate chromosomes. To increase the specificity of copy number inference, we manually generated a genomic exclusion list for the sex chromosomes. This list includes the centromeres, regions of low mappability (mappability score within 1 kb is less than 0.5[14]), regions with extremely high read count signals, and regions in Y chromosome with high signal in female samples. With these regions excluded, we called copy number variants with the FACETs algorithm. Confirming the validity of our method, we obtained single copy estimates for X and Y in all male cases, with diploid calls for PAR1 and PAR2 (**Supplementary Figure S3**). Identified CNVs were checked manually for accuracy.

**Mutation recurrence analysis**

Due to the extremely low number of mutations, we manually interrogated selected genomic elements for the presence of more than one mutation. These elements included coding regions of genes (exons), transcription start sites[15], 3'UTRs and 5'UTRs (downloaded from UCSC). In addition, we included regulatory elements in the *FIP1L1-PDGFRA* positive EOL1 leukemia cell line: putative active promoters and enhancers were inferred from H3K27ac ChIP-Seq and insulators from CTCF ChIP-seq[16]. MACS2[17] was used for signal peak calling, and overlap with somatic mutations was calculated with BedTools[12]. Each ChIP-seq peak was considered a region of interest.

**Phylogenetic inference**

Inference of clonal structure of single tumors was performed with PhylogicNDT[18] (https://github.com/broadinstitute/PhylogicNDT) with ABSOLUTE output files and default parameters.

**Analysis of clonal hematopoiesis (CH)**

Somatic mutation calling for SNVs, insertions and deletions was performed on normal control samples using MuTect[7] and MuTect2[8] in single-sample mode. Variant calling was restricted to genomic loci of previously reported CH genes[19]. Only variants passing internal filters were selected, and germline variants were removed by comparison with matched HaplotypeCaller results using Tabix (version 0.2.6[20]) and BCFtools (version 1.8[21]). Oncotator[22] version 1.9.0.0 with Gencode v19 was used for annotation.

**Data sharing statement**

Whole genome sequences, germline and somatic variants will be available with access control via dbGAP.


1 Klion, A. D. *et al.* Familial eosinophilia: a benign disorder? *Blood* **103**, 4050-4055, doi:10.1182/blood-2003-11-3850 (2004).

2 Cibulskis, K. *et al.* ContEst: estimating cross-contamination of human samples in next-generation sequencing data. *Bioinformatics* **27**, 2601-2602, doi:10.1093/bioinformatics/btr446 (2011).

3 Costello, M. *et al.* Discovery and characterization of artifactual mutations in deep coverage targeted capture sequencing data due to oxidative DNA damage during sample preparation. *Nucleic acids research* **41**, e67, doi:10.1093/nar/gks1443 (2013).

4 Chen, X. *et al.* Manta: rapid detection of structural variants and indels for germline and cancer sequencing applications. *Bioinformatics* **32**, 1220-1222, doi:10.1093/bioinformatics/btv710 (2016).

5 Morton, L. M. *et al.* Radiation-related genomic profile of papillary thyroid carcinoma after the Chernobyl accident. *Science* **372**, doi:10.1126/science.abg2538 (2021).

6 Shen, R. & Seshan, V. E. FACETS: allele-specific copy number and clonal heterogeneity analysis tool for high-throughput DNA sequencing. *Nucleic acids research* **44**, e131, doi:10.1093/nar/gkw520 (2016).

7 Cibulskis, K. *et al.* Sensitive detection of somatic point mutations in impure and heterogeneous cancer samples. *Nature biotechnology* **31**, 213-219, doi:10.1038/nbt.2514 (2013).

8 Auwera, G. v. d. & O'Connor, B. D. *Genomics in the cloud : using Docker, GATK, and WDL in Terra*. First edition. edn, (O'Reilly Media, 2020).

9 Saunders, C. T. *et al.* Strelka: accurate somatic small-variant calling from sequenced tumor-normal sample pairs. *Bioinformatics* **28**, 1811-1817, doi:10.1093/bioinformatics/bts271 (2012).

10 Carter, S. L. *et al.* Absolute quantification of somatic DNA alterations in human cancer. *Nature biotechnology* **30**, 413-421, doi:10.1038/nbt.2203 (2012).

11 Kasar, S. *et al.* Whole-genome sequencing reveals activation-induced cytidine deaminase signatures during indolent chronic lymphocytic leukaemia evolution. *Nature communications* **6**, 8866, doi:10.1038/ncomms9866 (2015).

12 Quinlan, A. R. & Hall, I. M. BEDTools: a flexible suite of utilities for comparing genomic features. *Bioinformatics* **26**, 841-842, doi:10.1093/bioinformatics/btq033 (2010).

13 Mermel, C. H. *et al.* GISTIC2.0 facilitates sensitive and confident localization of the targets of focal somatic copy-number alteration in human cancers. *Genome Biol* **12**, R41, doi:10.1186/gb-2011-12-4-r41 (2011).

14 Derrien, T. *et al.* Fast computation and applications of genome mappability. *PloS one* **7**, e30377, doi:10.1371/journal.pone.0030377 (2012).

15 Rheinbay, E. *et al.* Recurrent and functional regulatory mutations in breast cancer. *Nature* **547**, 55-60, doi:10.1038/nature22992 (2017).

16 Consortium, E. P. *et al.* Expanded encyclopaedias of DNA elements in the human and mouse genomes. *Nature* **583**, 699-710, doi:10.1038/s41586-020-2493-4 (2020).

17 Zhang, Y. *et al.* Model-based analysis of ChIP-Seq (MACS). *Genome Biol* **9**, R137, doi:10.1186/gb-2008-9-9-r137 (2008).

18 Leshchiner, I. *et al.* Comprehensive analysis of tumour initiation, spatial and temporal progression under multiple lines of treatment. *bioRxiv*, 508127, doi:10.1101/508127 (2019).

19 Jaiswal, S. *et al.* Age-related clonal hematopoiesis associated with adverse outcomes. *The New England journal of medicine* **371**, 2488-2498, doi:10.1056/NEJMoa1408617 (2014).

20 Li, H. Tabix: fast retrieval of sequence features from generic TAB-delimited files. *Bioinformatics* **27**, 718-719, doi:10.1093/bioinformatics/btq671 (2011).

21 Li, H. & Durbin, R. Fast and accurate short read alignment with Burrows-Wheeler transform. *Bioinformatics* **25**, 1754-1760, doi:10.1093/bioinformatics/btp324 (2009).

22 Ramos, A. H. *et al.* Oncotator: cancer variant annotation tool. *Human mutation* **36**, E2423-2429, doi:10.1002/humu.22771 (2015).

#### Supplementary Figure Legends

**Supplementary Figure S1. A,** Time between first and second (remission) draw for all patients. **B,** Mean sequencing depth for tumor (blue) and normal (orange) samples. **C,** Detailed view of the genomic breakpoints in participating genes *PDGFRB*, *IQGAP2* and *UVRAG.* Split sequencing read portions are highlighted with color. Other reads are shown in gray. Total read depth is shown on top, with reduced depth indicative of deleted sequence evident between breakpoints.

**Supplementary Figure S2: A,** Variant allele fraction distribution for SNVs, insertions and deletions after filtering. **B,** Mutation spectrum of a second signature identified from SNVs in this cohort. **C,** Sum of probabilities to belong to one of the two signatures for each mutation and patient. **D**, Variant allele fraction for each mutation depending on the probability to belong to signature S2-SBS43. **E,F,** Individual mutation spectra for all SNV mutations given a three-base sequence context before (**E**) and after (**F**) signature-based filtering. Colors indicate base changes (reference base to mutated base). Position on the grid refers to the reference sequence context around the mutated base.

**Supplementary Figure S3.** Distribution of sequencing depth for different genomic element types.

**Supplementary Figure S4.** Copy number estimation for male sample HES2 (**A**) and female sample HES15 (**B**) showing the log2 copy ratio for tumor and normal (top), the log-odds ratio for heterozygous single-nucleotide variants (second row) and estimated total copy number for each chromosome (bottom). Each heterozygous SNP is represented by a light blue dot. Red lines in the top and middle panel indicate called segments. Black bar in the bottom panel shows the total copy number, with copy number of the minor allele in red. In the male sample, the total copy number of X and Y, respectively is 1. The copy number of the PAR regions is 2 (one from X, one from Y), with copy number one of both the major and minor alleles. In the female sample, the total copy number of X is 2, and Y is zero.

#### Supplementary Tables

**Table S1. Participant information**

**Table S2. Sequencing metrics**

### Figure S1

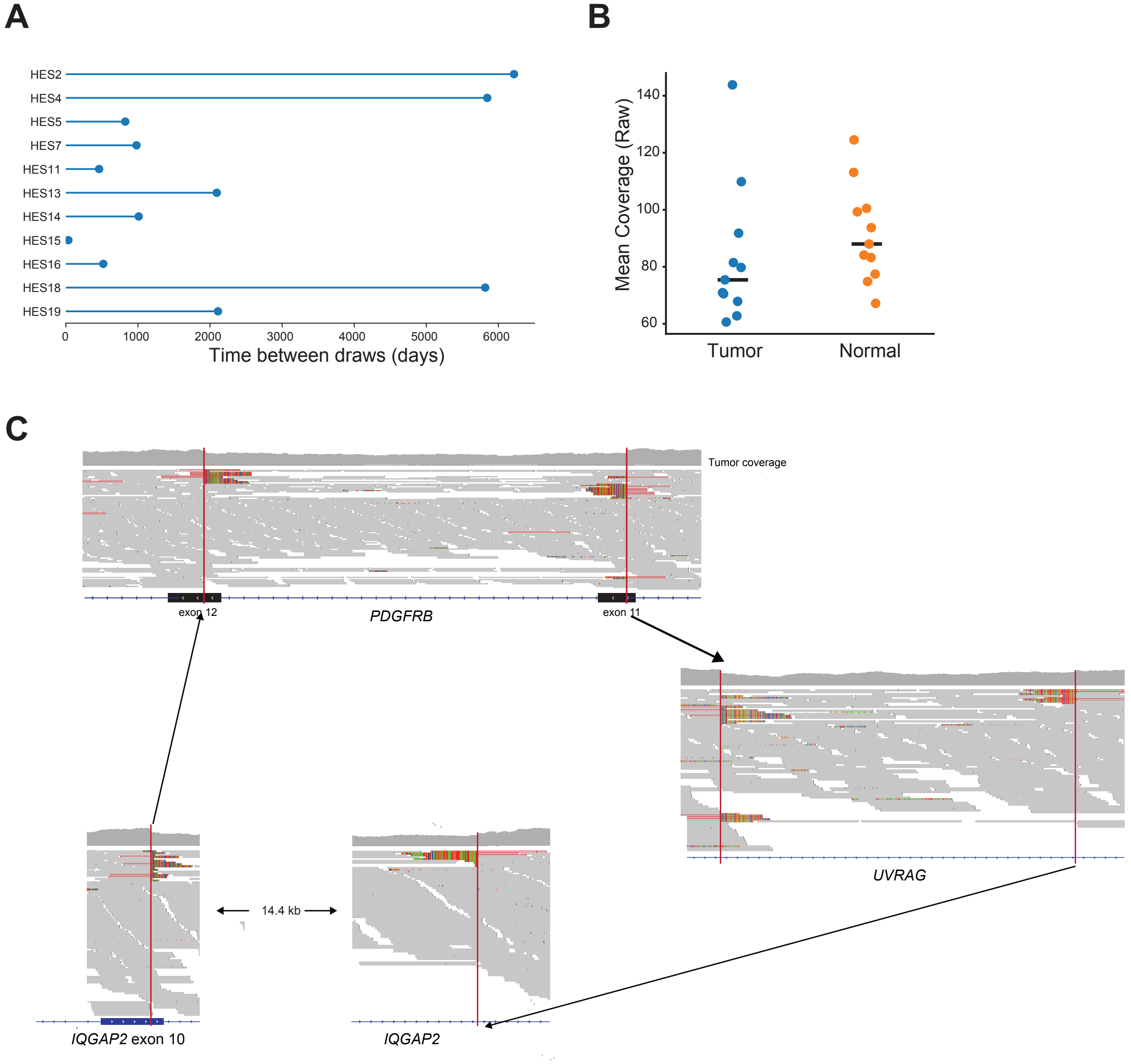
A
HES2
HES4
HES5
HES7
HES11
HES13
HES14
HES15
HES16
HES18
HES19
0
1000
2000
3000
4000
5000
6000
Time between draws (days)
B
Mean Coverage (Raw)
60
80
100
120
140
Tumor
Normal
C
Tumor coverage
exon 12
PDGFRB
exon 11
UVRAG
14.4 kb
IQGAP2 exon 10
IQGAP2

### Figure S2

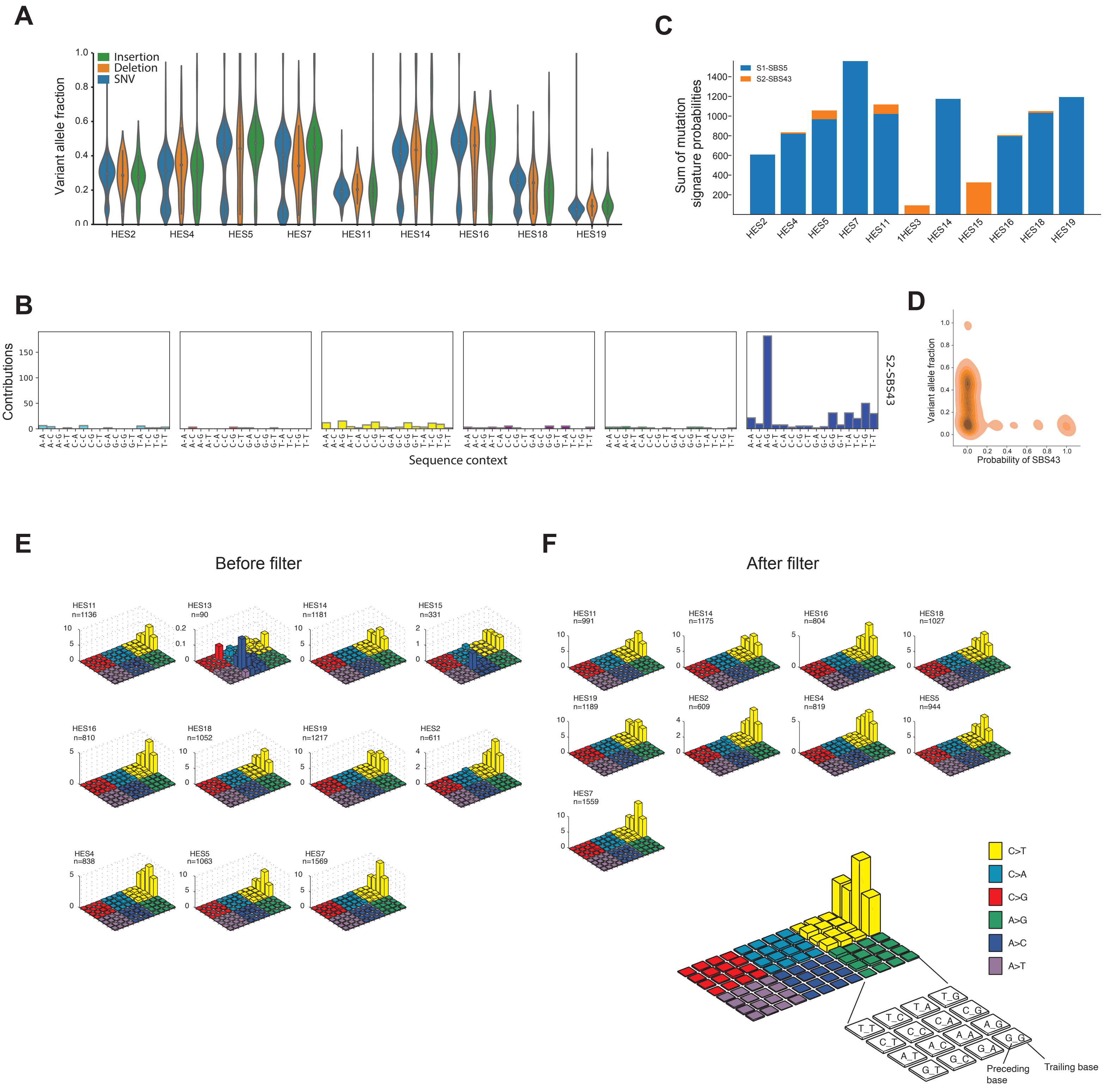

A
Insertion
Deletion
SNV
Variant allele fraction
HES2
HES4
HES5
HES7
HES11
HES14
HES16
HES18
HES19
C
S1-SBS5
S2-SBS43
Sum of mutation signature probabilities
B
Contributions
S2-SBS43
Sequence context
D
Variant allele fraction
Probability of SBS43
E
Before filter
F
After filter
C>T
C>A
C>G
A>G
A>C
A>T
Preceding base
Trailing base

### Figure S3

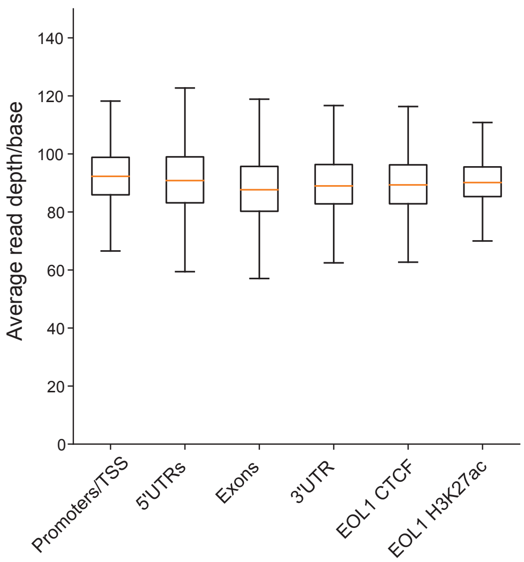

Average read depth/base
140
120
100
80
60
40
20
0
Promoters/TSS
5'UTRs
Exons
3'UTR
EOL1 CTCF
EOL1 H3K27ac

#### Figure S4

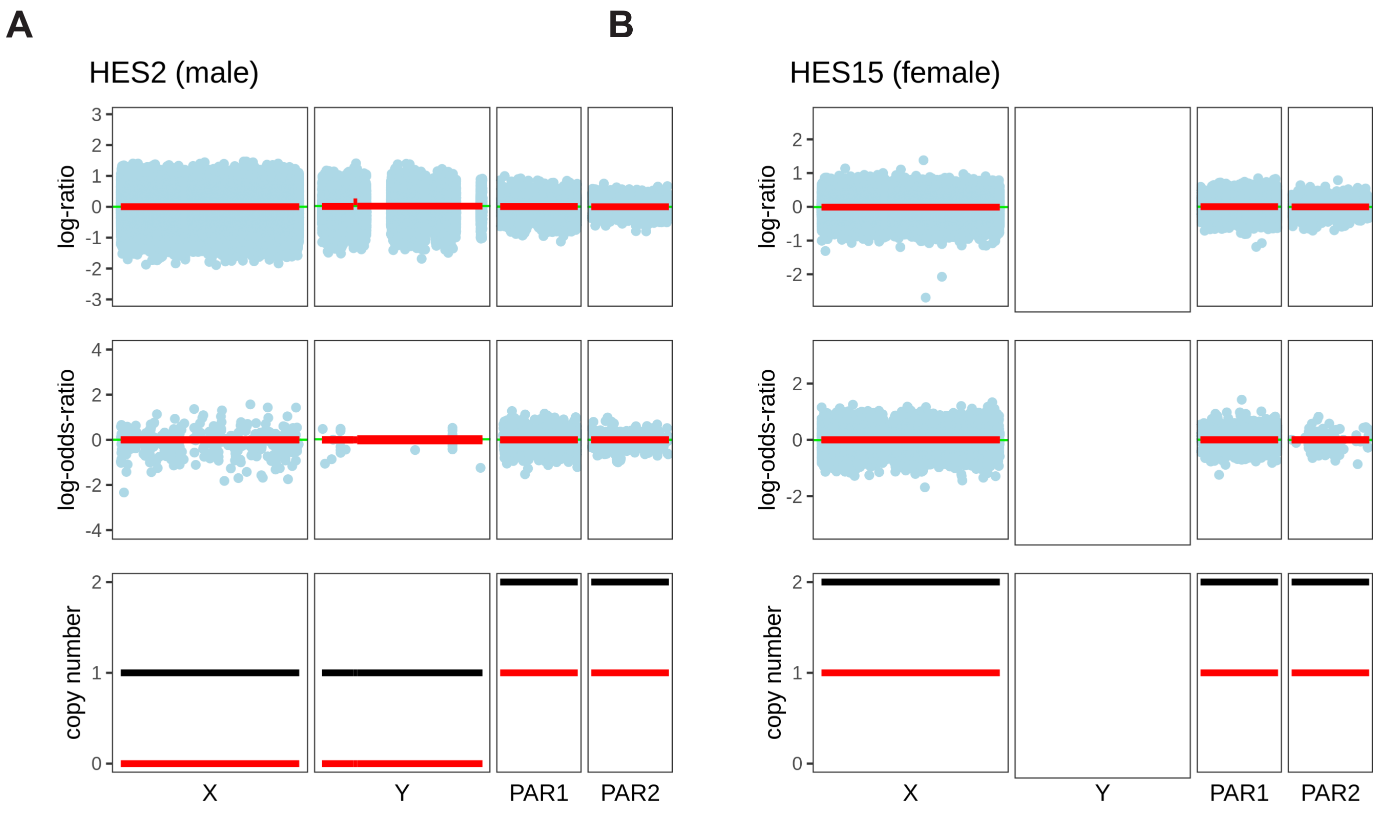
A
HES2 (male)
B
HES15 (female)
log-ratio
log-odds-ratio
copy number
X
Y
PAR1
PAR2
X
Y
PAR1
PAR2
